# Extinct and extant termites reveal the fidelity of behavior fossilization in amber

**DOI:** 10.1101/2023.05.22.541647

**Authors:** Nobuaki Mizumoto, Simon Hellemans, Michael S Engel, Thomas Bourguignon, Aleš Buček

**Affiliations:** Okinawa Institute of Science & Technology Graduate University, Onna-son, Okinawa, Japan; Division of Invertebrate Zoology, American Museum of Natural History, New York, NY, USA; Department of Ecology & Evolutionary Biology, University of Kansas, Lawrence, KS, USA; Division of Entomology, Natural History Museum, University of Kansas, Lawrence, KS, USA; Biology Centre, Czech Academy of Sciences, Ceske Budejovice, Czech Republic

**Keywords:** amber, collective behavior, fossil record, leadership, social interaction, movement coordination

## Abstract

Fossils encompassing multiple individuals provide rare direct evidence of behavioral interactions among extinct organisms. However, the fossilization process can alter the spatial relationship between individuals and hinder behavioral reconstruction. Here, we report a Baltic amber inclusion preserving a female-male pair of the extinct termite species *Electrotermes affinis*. The head-to-abdomen contact in the fossilized pair resembles the tandem courtship behavior of extant termites, although their parallel body alignment differs from the linear alignment typical of tandem runs. To solve this inconsistency, we simulated the first stage of amber formation, the immobilization of captured organisms, by exposing living termite tandems to sticky surfaces. We found that the posture of the fossilized pair matches trapped tandems and differs from untrapped tandems. Thus, the fossilized pair likely is a tandem running pair, representing the first direct evidence of the mating behavior of extinct termites. Furthermore, by comparing the positions of partners on a sticky surface and in the amber inclusion, we estimated to 67% the probability that the leader role in the fossilized tandem was performed by a male. Our results demonstrate that past behavioral interactions can be reconstructed despite the spatial distortion of body poses during fossilization. Our taphonomic approach clarifies how certain behaviors can be inferred from fossil occurrences.

## Introduction

Group-living animals coordinate their movement to stay together while searching for a safe place or feeding site (Couzin, 2018). Movement coordination is achieved by adjusting the speed and direction of movements in response to neighbors (Herbert-Read, 2016; Sumpter, 2006). Such behavioral interactions among group members affect the shape of the whole group (Hemelrijk and Hildenbrandt, 2012) and the spatial distribution among neighbors (Ballerini et al., 2008). Collective behavior can be observed in a diverse range of organisms, from bacteria to humans (Sumpter, 2006), indicating the ancient origin of movement coordination. However, evidence of behavioral coordination in the fossil record is rare (Hou et al., 2008). Movement coordination is a dynamic process leading to spatial structures that are often altered during fossilization (Boucot, 1996). Although several studies have attempted to infer the behavioral processes employed in the collective behavior of extinct animals (Hou et al., 2008; Mizumoto et al., 2019; Vannier et al., 2019), uncertainty remains in their interpretation.

To evaluate behaviors preserved in the fossil record, it is essential to determine the biotic and abiotic factors that may have influenced the preservation of extinct organisms. Amber, or fossil resin, provides uniquely detailed soft-body preservation with 3-dimensional features ideal for capturing snapshots of “frozen behaviors,” fossils with animals preserved in action (Boucot, 1996; Hsieh and Plotnick, 2020). The co-occurrence of multiple individuals (i.e., syninclusions) is a valuable source of information for tracking behavioral interactions between extinct organisms, including predator-prey relationships (Barden et al., 2020; Coty et al., 2014), mating behaviors (Fischer and Hörnig, 2019), and host-parasite associations (Dunlop et al., 2014; Jiang et al., 2021). Nevertheless, the preservation process in tree resin is not instantaneous and induces a range of behavioral responses that interfere with the behavior performed immediately before entrapment. Interaction with fresh sticky resin often results in loss of body parts, defecation, and induction of stress behavior, such as egg laying (Arillo, 2007). Thus, to improve the accuracy of inferring behavioral coordination from fossil occurrences, it is essential to identify how the behavior of animals is modified during entrapment and death.

Tandem-running behavior is the simplest movement coordination maintained by leader-follower behavioral interactions (Franks and Richardson, 2006; Valentini et al., 2020). In termites, tandem running is performed by a pair of unwinged female and male after the dispersal flight (Nutting, 1969). During the mating season, winged termites fly from their natal nests and disperse. After dispersal, both females and males land on the ground or tree trunks and run about searching for a mate. Once encountered, a pair forms a tandem run, in which one follows the other by maintaining close contact with the tip of the leader’s abdomen (Raina et al., 2003). The follower maintains contact with their antennae or mouthparts. Either females or males or both sexes can play the leader role, depending on the termite species (Mizumoto et al., 2022). The tandem pairs seek a suitable site to establish their nest and form a lifelong monogamous royal pair.

Eocene Baltic amber is historically the most productive deposit for Cenozoic fossiliferous resin and includes the most diverse fossil insect assemblages (Grimaldi and Engel, 2005), including examples of “frozen” behaviors (Fischer and Hörnig, 2019) as well as extinct and extant termite genera (Engel et al., 2007). Termites are abundant in the Baltic amber, particularly winged and de-winged termite reproductives. Tandem pairs walking on or near tree trunks (Mizumoto et al., 2020) are susceptible to entrapment in tree resin, yet fossils of tandem pairs have never been described. Here, we report a unique fossil occurrence of syninclusions of two dealate termite imagoes of *Electrotermes affinis* in 38-million-year-old Baltic amber (Fig. 1). The syninclusions portray a heterosexual composition of the pair, head-to-abdomen orientation, without wings, which are consistent with the two termite individuals representing the first and earliest documented occurrence of a fossil tandem pair. However, the side-to-side alignment of the termite pair distinguishes it from the stereotypical front-back alignment in natural tandem runs. We hypothesized that this spatial organization formed during the process of entrapment in the resin and we tested this hypothesis by empirically simulating the entrapment process on living termite mating pairs using sticky traps. This taphonomic experiment enabled quantitative measures for more reliably documenting “frozen” behaviors preserved in amber.

**Figure 1.**
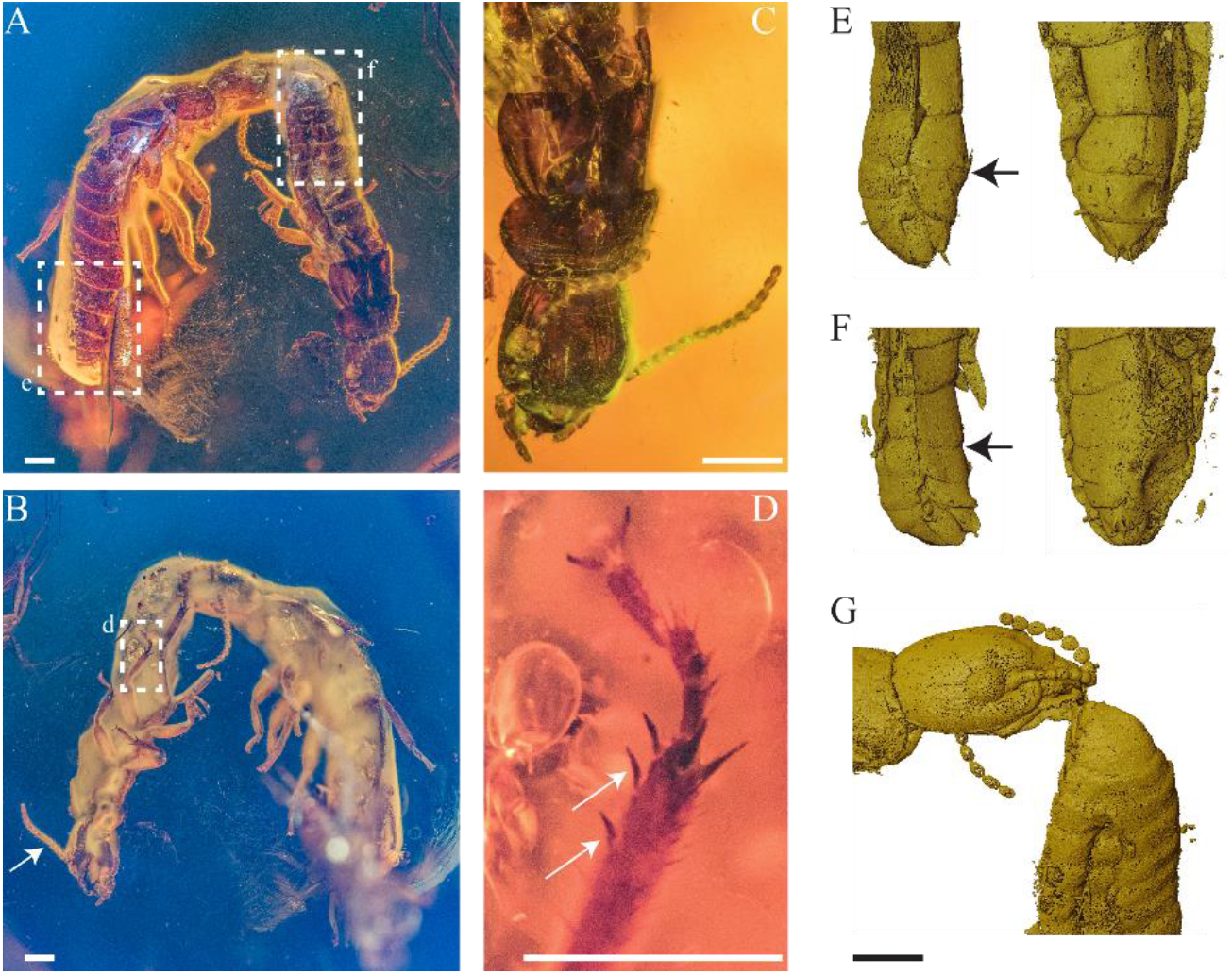
*Electrotermes affinis* pair in Baltic amber. (A-B) Habitus view of the dorsal and ventral sides of the tandem, respectively, with (B) an arrow pointing to the 15-articles antenna of the tandem leader. (C) Focus featuring the absence of fontanelle and the pronotum being broader than the head, a combination of features typical of Kalotermitidae. (D) Focus on the middle tibia bearing the two outer spines (see arrows) characteristics of the fossil genus *Electrotermes*. (E-G) X-ray uCT scans of the abdomen tips of the tandem (E) follower and (F) leader, with the (G) point-of-contact between the follower’s head and the leader’s abdomen. Sternite anatomy (arrows indicate the upper limit of the seventh sternite) revealing that the (F) tandem leader is male, and the (E) follower is female. Scale bars are 0.5 mm.

## Results

### Amber syninclusion of de-winged female and male termites

The Baltic amber syninclusion contains two de-winged drywood termite imagoes (Kalotermitidae) (specimen NMP T3532, Fig. 1A). An opaque cloud of bubbles visually obscured the ventral posterior parts of the abdomens of the two termites. X-ray microtomography uncovered that one individual was female (Fig. 1AE) and the other was male (Fig. 1AF), based on the anatomy of the 7th sternite, which is enlarged only in females (Weesner, 1969). Spines on the mesotibia indicated that both termites belong to the extinct genus *Electrotermes* (Krishna, 1961) (Fig. 1BD). Body lengths of ∼6.5 mm for the female and ∼5.5 mm for the male further confirmed that the species is *Electrotermes affinis* rather than the distinctly smaller *Electrotermes girardi* (Engel et al., 2007; Weidner, 1955). The larger body size of the female imago is consistent with the sexual dimorphism of extant species of Kalotermitidae (Afzal, 1984; Miyaguni et al., 2021), and further corroborates the sex determinations. The female’s mouthparts were in contact with the tip of the male’s abdomen (Fig. 1G), implying that these two termites were performing tandem-running behavior at the moment of entrapment. However, the female and the male within the amber piece were positioned side-by-side (Fig. 1AB), which is unlike living termites in which the leader and follower are in a single file, with the follower just behind the leader (Fig. 2A). We hypothesized that the non-instantaneous fixation of trapped termites by sticky tree resin is at the origin of their parallel orientation preserved in the amber.

**Figure 2.**
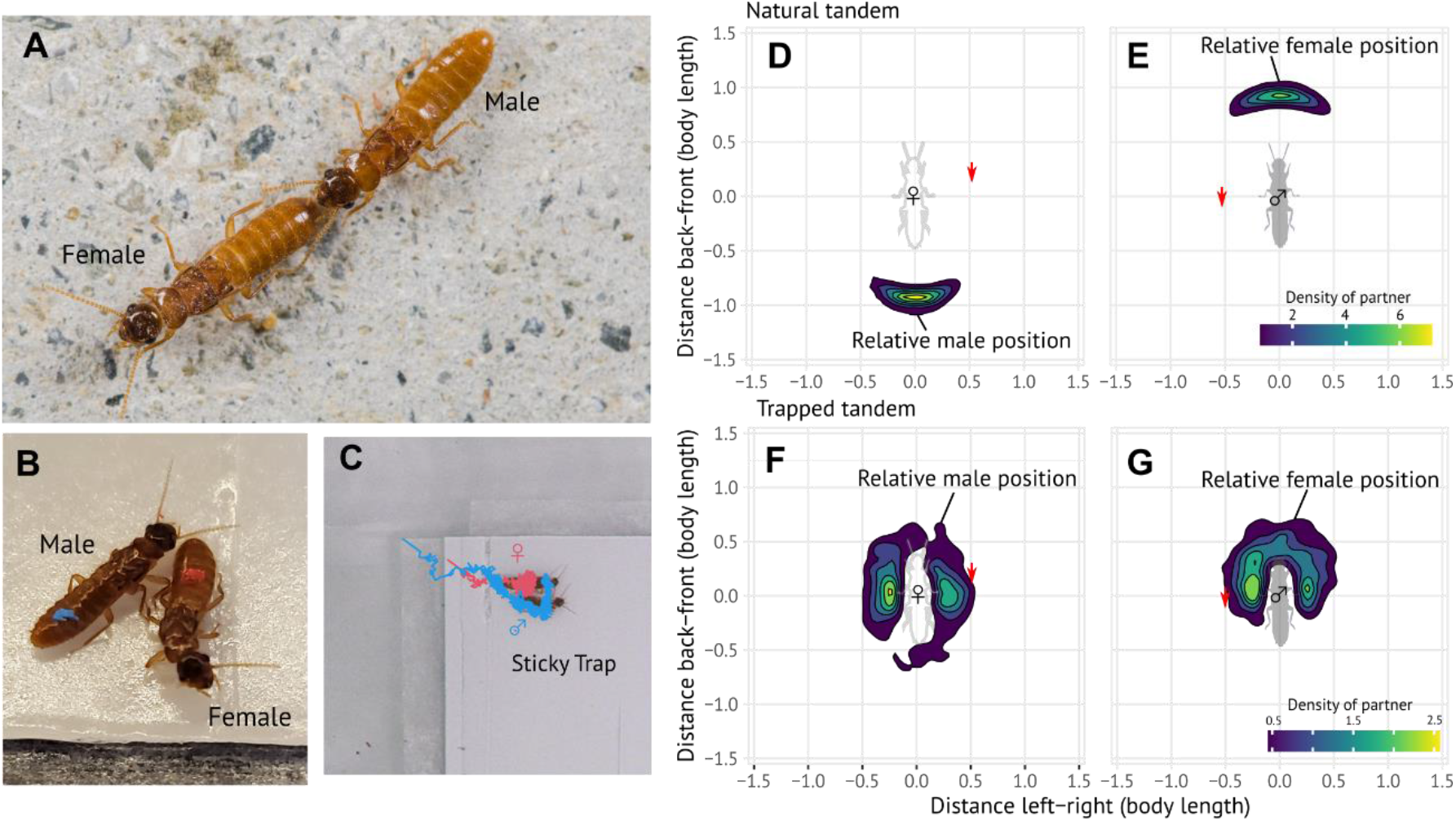
The relative position of females and males forming mating pairs. (A-C) Mating pairs of the termite *Coptotermes formosanus* in (A) a regular tandem run and (B-C) on a sticky surface. Females are marked in red, and males in blue. The convoluted lines indicate the trajectories of a female and a male during the 30 minutes after the pair entered the sticky trap. (D-G) Density map showing the position of one partner, represented by the central shadow of a termite heading towards the top, in relation to the other. The density map was based on pooled data of (D-E) natural tandem pairs and (F-G) tandem pairs trapped on the sticky surface. Red arrows indicate the data point of the amber inclusion.

### Phases of simulated entrapment and movement dynamics

To evaluate the effect of the entrapment process on tandem-running behavior, we observed movement patterns of termite mating pairs on a sticky surface. We simulated the tree resin using a sticky trap for insect collection, a method that has been used to mimic the process of resin sampling (Solórzano Kraemer et al., 2018, 2015). We then compared the fossil information with the spatial organization of the leader-follower relationships during natural tandem runs (Fig. 2A) and after being trapped by sticky traps (Fig. 2B).

We introduced a mating pair of the termite *C. formosanus* into an experimental arena with a sticky trap (Fig. S1) and recorded the events during which termite leaders entered the sticky surface (*n* = 26). Although the trap partially obstructed the movement of leaders that naturally entered the sticky traps before the follower, followers did not stop following the leader, resulting in both individuals entering completely, with the whole body area and all appendages onto the sticky surface for all events (Video S1). On the sticky surface, the movement speed was significantly lower than in natural tandems (sex pooled, Mean ± SD, natural: 1.99 ± 0.51 body-length/sec, sticky: 0.06 ± 0.02 body-length/sec; Wilcoxon rank sum test, *P* < 0.001). Although some individuals escaped from the surface, both the female and male were trapped in 17/26 events after 10 min, 14/26 events after 20 min, and 9/24 events after 30 min. There was no significant difference in the probability of escaping the sticky surface between sexes (mixed effect Cox model, χ^2^_1_ = 0.214, *P* = 0.644). Thus, our sticky trap successfully simulated the situation where termites were not instantaneously immobilized but could behaviorally respond to the entrapment (Fig. 2C), potentially resulting in the specific spatial orientation observed in the amber inclusion (Fig. 1).

### Spatial organization of trapped pairs

The spatial orientation of the leader and the follower after entrapment was significantly different than in natural tandem runs. The distance between the body centroids of the leader and the follower was smaller in trapped pairs than in natural tandems (Fig. 2D-G, Fig. S2, Exact Wilcoxon rank sum test, *W* = 599, *P* < 0.001). This is because partners of trapped pairs were often positioned side-by-side, differing from the linear positioning of natural tandems (Fig. 2D-G). Surrogate distances artificially generated by randomly pairing leaders and followers from different trapped tandems showed larger inter-individual distances than in observed trapped tandems (Fig. S2), suggesting that the smaller distance observed on the sticky trap was the result of interaction between the leader and the follower. Furthermore, partners of trapped pairs were either heading in the same or opposite direction, unlike partners of surrogate tandems (Fig. S3, the distributions of heading direction difference was different among conditions; KS test, *P* < 0.001). Thus, the process of becoming ensnared on a fresh resin surface modifies the spatial structures of tandem runs in a unique and specifiable way. The observed interindividual distances in the amber inclusion fell well within the range of trapped tandem pairs and outside that of natural tandem runs (Fig. 2, Figs S2). Therefore, the observed side-by-side positioning of the two individuals of *E. affinis* in the amber inclusion is expected for an entrapped tandem run.

### Estimating leader-follower roles from relative body postures

We estimated the leader-follower roles of the two individuals of *E. affinis* in the amber inclusion by comparing them with the observations of trapped *C. formosanus*. For comparison purposes, we created surrogate datasets by artificially swapping the sex identities of *C. formosanus*. Because the female is always the leader in *C. formosanus*, the male is always the leader in the surrogate datasets. We estimated the probability for each tandem member to be the leader or follower by comparing the relative posture of the fossilized tandem to the distribution of relative postures observed in the experimentally trapped tandems and the surrogate datasets.

To quantify the relative postures of termites, we extracted the coordinates of body parts (head, pronotum, and abdominal tip for each sex). We measured 15 pairwise distances representing all possible combinations of pairwise distances between the three body parts of the two partners. We then performed a principal component analysis to reduce the dimensionality of the datasets (Fig. 3, see Table S1 for principal component loadings).

**Figure 3.**
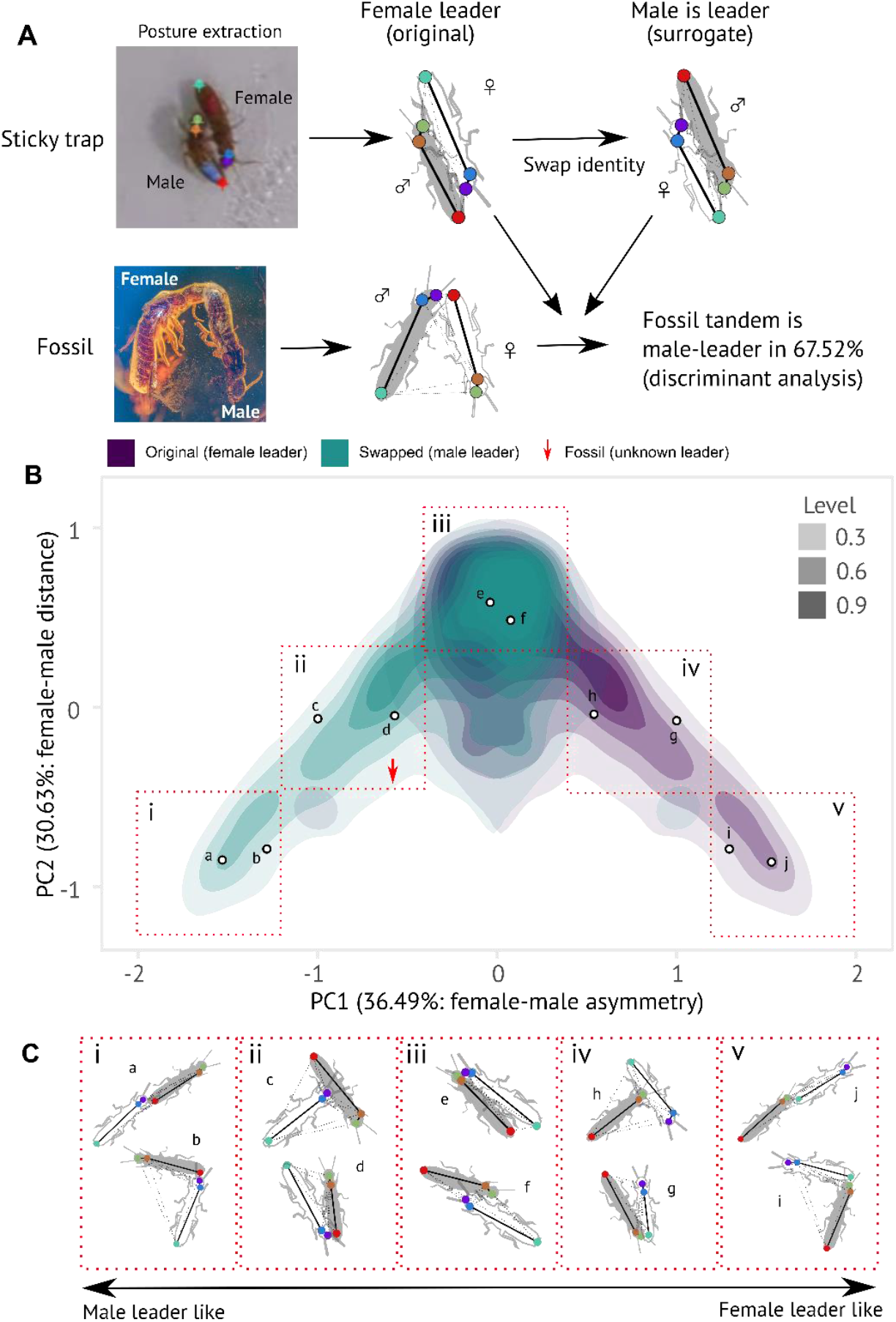
Inference of leader-follower roles from relative postures of termite pairs trapped by sticky surface. (A) Analysis procedure. For each tandem pair, we extracted the body part coordinates (middle of the head, pronotum, and abdomen tip) of both partners and calculated all 15 pairwise distances. We created the surrogate datasets by swapping the identities of females and males. As the female is always the leader in *C. formosanus* tandems, the male is always the leader in the surrogate datasets. We performed a principal component analysis (PCA) and a discriminant analysis (DA) with the first four components for all datasets. (B) Scatter plot of the PCA based on original and surrogate datasets. The red arrow indicates the position of the two individuals of *E. affinis* in the amber inclusion. (C) Postures of representative tandem pair data points.

The first four principal components explained 97% of the observed variance (PC1: 36.5%, PC2: 30.6%, PC3: 16.5%, PC4: 13.5%). There was no separation of relative postures between the original and surrogate datasets along PC2 to PC4 (Fig. S4). Thus, PC2-4 reflected the relative posture of the tandem irrespective of the sex identity of the leader and follower. In contrast, the relative postures of the original and surrogate datasets were separated along the PC1, where the data points of the original and surrogate datasets formed mirror images (Fig. 3B, S4). Relative postures with the characteristic male-led tandems had negative values along the PC1 axis, while female-led tandem postures had positive values. In the regions along the PC1 that departed the most from zero, the leader and follower roles can be assigned with high confidence based on the relative posture of the partners (regions *i* and *v* in Fig. 3B, respectively). The relative posture of untrapped tandems also occupied these regions (Fig. S5). The intermediate regions (regions *ii* and *iv* in Fig. 3B) included tandem runs for which the leader and follower roles could only be inferred with low confidence, not in a decisive manner, based on relative postures. In the central region of the PCA (region *iii* in Fig. 3B), the relative postures of the original and surrogate datasets were similar, and the partners of the tandem could not be assigned leader or follower roles. The fossilized pair of *E. affinis* occupied a region of the PCA plot that mostly matched a male-leader-like posture (red arrow in Fig. 3B). Our linear discriminant analysis using PC1-4 assigned a 67.5% probability to a male-leading tandem for the fossil, and thus a 32.5% probability to the alternative scenario of a female-leading tandem.

## Discussion

Interacting organisms are preserved in amber inclusions in great detail. However, the fixation of organisms in the tree resin is not instantaneous and may alter behavior. Disentangling the natural behavior related to social interactions from the behavioral responses to entrapment is a major challenge in the analysis of behavior fossilized in amber. Combining a fossil record with taphonomical behavioral experiments, our results provide direct evidence that a kalotermitid termite, *E. affinis*, performed tandem movement coordination 38 million years ago. This interpretation is supported by the ancestral state reconstruction of tandem running behavior in termites, which supported that tandem running behavior was present in the common ancestor of modern termites, and has been preserved across kalotermitid species (Mizumoto et al., 2022). Therefore, the tandem running behavior of *E. affinis* is the oldest record of a movement coordination behavior still present across extant relatives. Importantly, our empirical simulations with extant termites indicate that the spatial distribution of the tandem pair partners found in amber is expected to differ from that of untrapped tandems, while still retaining spatial patterns allowing their identification as tandem pairs. Based on the spatial orientation of the interacting termites in amber reported herein, our analyses suggest that the male is more likely the tandem leader. This is consistent with the inferred ability of the common ancestor of extant Kalotermitidae to form both male-led and female-led tandems (Mizumoto et al., 2022). The uncertainty of such role assignment is, however, high due to the disturbance of tandem behavior during entrapment. By simulating the process of living animals being entrapped by sticky objects, our study quantitatively clarified what aspects of behaviors can and cannot be inferred from fossil records.

Some fossils preserve the “frozen” behavior of animals in actions at the moment of death (Boucot, 1996; Hsieh and Plotnick, 2020). However, our results demonstrate that animals on the sticky trap are not instantaneously immobilized and change their postures on the surface. These experiments imply that the spatial orientation of animals preserved in sticky matrices, such as in tree resin prior to fossilization into amber, is influenced by the process of entrapment. Therefore, the interpretation of fossilized behavior can be dramatically refined or even corrected by observing the behavior of living organisms under entrapment conditions. Some behaviors fossilized in amber may remain unaltered by the entrapment process. For example, the preservation of mating moths in *copula* (Fischer and Hörnig, 2019) or hell ants grasping prey items (Barden et al., 2020) suggests that the inter-individual interactions of these behaviors are strong enough not to be disturbed by the movement on the sticky surface. However, entrapment in amber likely affects many other behaviors. For example, insects dispersing through phoresy can be preserved detached from the host insect, perhaps because the host struggled on the sticky surface before complete encasement (Robin et al., 2019). The consequence of different behavioral responses can be studied using extant relatives. Furthermore, animals have evolved behavioral responses to sticky objects. For example, recent studies have revealed that ants are not passively affected by sticky objects but actively modify them. Red imported fire ants cover sticky surfaces with soil particles to access food resources (Wen et al., 2021), and granivorous desert ants remove sticky spider webs from nestmates to rescue them (Kwapich and Hölldobler, 2019). Scavenging insects can be attracted by large animals trapped on a sticky surface (Solórzano Kraemer et al., 2018), and the spatial distribution of these insects may have reflected their foraging behavior. Thus, future studies on behavioral responses to sticky objects by animals will increase our understanding of fossil records in amber, as well as shed light on the behavioral capacity of extant insects.

Interestingly, the disturbance of tandem pair movements by a sticky surface is qualitatively distinctive from other sources of disturbance in termite mating pairs. For example, when a tandem pair encounters a predatory ant that captures one of the two partners, an escape behavior is triggered in the other individual (Li et al., 2013; Matsuura et al., 2002). When the tandem pair is interrupted by other termites or separated spontaneously, the leader pauses, and the follower moves to search for a partner (Mizumoto et al., 2020; Mizumoto and Dobata, 2019). Hence, termite mating pairs react dynamically to external disturbances in these situations. In contrast, when a sticky trap caught the leader, the follower neither escaped nor left the leader to search for an alternative partner. Instead, the follower walked around the leader and was caught by the sticky trap, resulting in both partners being immobilized side-by-side (Fig. 4). Therefore, sticky surfaces can catch groups of animals, one after another. Social insects, such as ants and termites, are frequently found in groups as amber inclusions (Coty et al., 2014; Zhao et al., 2020). Our results show that social interactions of extinct organisms can be inferred from these amber syninclusions using a quantitative approach measuring the posture and orientation of fossils in amber, with a comparison of behavioral observations of extant relatives.

**Figure 4.**
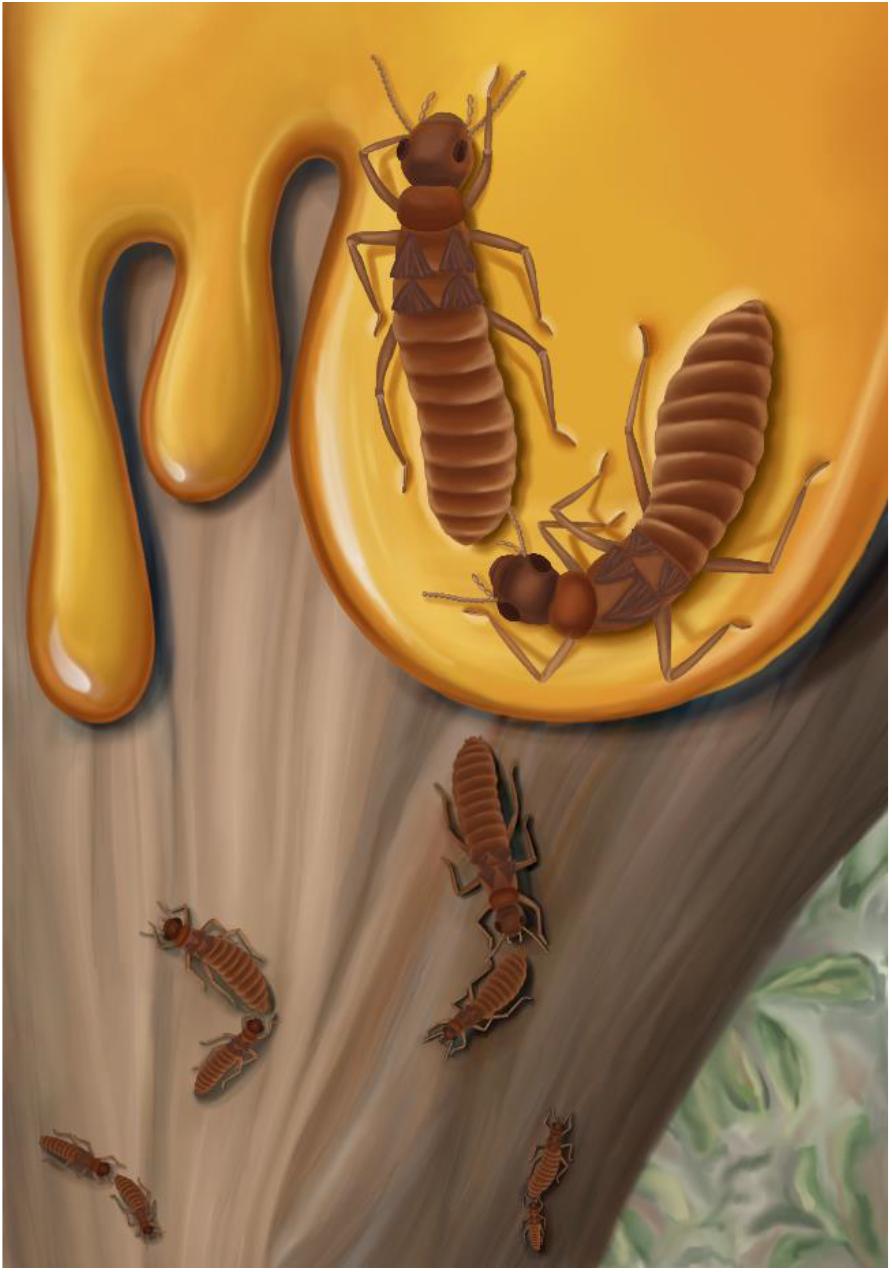
Artistic reconstruction of *Electrotermes affinis* tandem pairs running freely on a tree bark and one tandem trapped by tree-resin.

In conclusion, we show that spatial data obtained from behavioral observations of animals on a sticky surface are suitable for inferring the posture and relative positioning of animals expressing specific behavior at the time of ensnarement in amber. The behavioral responses of animals to the sticky surface may be slightly different when the sticky surface is not solid but liquid and viscous. The viscosity of resins could be variable depending on the local conditions, age of maturation, and tree species (Solórzano Kraemer et al., 2018). Future studies are needed to investigate these preservation factors. Using an integrative approach to simulate the process of becoming ensnared in resin, fossil records in amber have an untapped potential to uncover the behavior repertoire of extinct animals, especially their social interactions.

## Materials and Methods

### Microtomography of amber inclusion

The amber inclusion containing one female and one male termite originates from the Yantarny mine, Kaliningrad (Russia). It is a polished piece 40 mm long, 23 mm wide, and 7 mm high, and its weight is 4.40 g (see Fig. S6 for overall fossil habitus). The amber fossil was deposited at the National Museum, Prague (Czechia) as specimen NMP T3532. We performed µCT scans of the amber inclusion using a Zeiss Xradia 510 Versa 3D X-ray microscope and the Zeiss Scout-and-Scan Control System software (v11.1.6411.17883). A total of 1601 projections were collected during 360° rotation of the sample mounted on a rotating carousel. Three scans were performed: two scans of the abdominal tip regions of both termite individuals and one overall scan including both individuals. For the overview of scanning parameters, see Table S2. The 3D reconstructions of collected projections were performed with the Zeiss Scout-and-Scan Control System Reconstructor software (v11.1.6411.17883). Scan datasets were visualized with the Amira software (v6.7). The voxels representing termite specimens were selected using the “Threshold “function, and subsequently, the outer termite morphology was visualized using the “Volume Rendering “function of the Amira software. Rotation animation was rendered using the “Animation “module.

### Behavioral observations

Alates of *C. formosanus* were collected using light trapping in Okinawa, Japan, between May and June 2021. After collection, we brought them back to the lab and used individuals that shed their wings for behavioral observations. Individuals were separated by sex and kept on the moistened filter paper until the experiments. All observations were made within 12 hours after collection.

We simulated the amber entrapment process using sticky traps for insect collection. Sticky traps have been used to estimate the sampling bias of the insect assemblages captured in amber in tropical forests (Solórzano Kraemer et al., 2018, 2015). The processes of entrapment in tree resin have notable differences from entrapment on sticky tapes, such as different adhesive properties and the engulfing properties of fresh tree resin absent in sticky tape. However, arthropod assemblages found in amber or tree resin are similar to those captured by sticky traps (Solórzano Kraemer et al., 2018, 2015), suggesting that sticky traps mimic tree resin adequately. We prepared an experimental arena by attaching a trimmed sticky trap (square with 80 mm side; 2-7362-01, ASONE, Japan) to the center of a plastic container (221 × 114 × 37 mm) with double-sided tape (Fig. S1). The surface of the plastic container was polished so that termites could walk on the surface without slipping. We introduced a tandem running female-male pair of termites in the arena. The pair was covered by a Petri dish (Φ=40mm) until both partners restarted the tandem run (Fig. S1). We recorded the behavior of every pair for at least 20 minutes after they entered the sticky trap with a video camera (HC-X1500-K, Panasonic) at 30 FPS. We stopped observations for pairs that did not enter the sticky trap for 180 minutes (*n* = 8). Some pairs entered the trap and successfully escaped from the trap. In this case, we resumed the observation until they re-entered the trap. In total, we obtained 26 trap-entering events from 22 pairs. We compared the probability of escaping the trap between sexes, using the mixed-effects Cox model (coxme() function implemented in the R package coxme (Therneau, 2015)), with each pair ID included as a random effect.

We used the dataset of natural tandem runs in *C. formosanus* generated by (Mizumoto and Bourguignon, 2022) as a control. The experiments were performed using individuals collected on the same day and in the same manner as for the entrapment experiment. The experimental arenas consisted of a petri dish (φ = 90 mm) filled with moistened plaster. The surface of the arena was cleaned by rubbing off a thin layer of plaster before each trial. Arenas were placed in an acrylic cube box (200 mm) over which a Raspberry Pi Camera Module was mounted on the top board. Camera modules were connected to a Raspberry Pi 4 Computer Model B, and video was recorded using RPi-Cam-Web-Interface (https://elinux.org/RPi-Cam-Web-Interface) at 25 FPS. 30-minute video recordings from 27 pairs were available.

### Spatial organization

We extracted the coordinates (body centroid) and estimated the heading direction of termites from all videos using the video-tracking system UMATracker (Yamanaka and Takeuchi, 2018) (See Fig. S7 for all trajectories). We downsampled all videos to one frame per second (FPS) for the downstream analyses. From the data of coordinates and heading directions, we calculated the distance between a female and a male (in body length) and the position of each partner relative to the heading direction of its focal partner (relative direction, [0, π]). The relative direction ranged from 0 (the partner was just in front of the focal individual) to π (the partner was just behind the focal individual). We compared the mean values of these parameters between natural tandem runs and trapped tandem pairs using the linear mixed effect model (LMM) with the date of collection as random effect. For these analyses, we only used a subset of videos frames in which a female and a male were within a distance of twice a termite body length (calculated by summing up female and male body length for each video). This is a conservative threshold for the maximum distance at which two termites forming a tandem can interact (Valentini et al., 2020). The likelihood ratio test was used to determine the statistical significance of the explanatory variable (type II test).

To ensure that the spatial organization observed on sticky surfaces results from inter-individual interactions between partners and not artefacts that may be due to immobilization, we compared the parameters measured from the experimental datasets with surrogate datasets. Surrogate datasets were artificially created by pairing coordinates of females and males belonging to different tandem pairs after adjusting initial coordinates and the heading direction of swapped males to that of the original males. This randomization process breaks possible behavioral interactions within the surrogate pair, while maintaining the influence of a sticky surface on both females and males from different pairs (Valentini et al., 2020). We repeated this randomization process 1,000 times, obtaining 1,000 surrogate datasets. We measured the same parameters as above to compare the results with the original datasets.

### Posture analysis

We used DeepLabCut (version 2.2.1.1) for body part tracking (Mathis et al., 2018; Nath et al., 2019). Specifically, we labeled 194 frames taken from 25 videos (95% of which was used for training). We used a ResNet-50 (He et al., 2015; Insafutdinov et al., 2016) neural network with default parameters for 750,000 training iterations. We used one shuffle for validation and found that the test error was: 2.37 pixels, train: 1.07 pixels (image size was 164-640 by 164-412 pixels). We used a p-cutoff of 0.9 to condition the X-Y coordinates for downstream analyses. This network was then used to analyze videos. We tracked six body parts: female head, female pronotum, female abdomen, male head, male pronotum, and male abdomen, where the head was consistently labeled in the middle, the pronotum at the head-pronotum border, and the abdomen at the tip. We did not use multi-animal DeepLabCut (Lauer et al., 2022) but explicitly labeled female and male body parts separately. We calculated the 15 pairwise distances of the six different body parts for each frame.

Suppose the postures of the pairs on the sticky surface retain specific spatial properties reflecting their leader-follower role. In that case, one could infer the leader-follower role from the snapshot observed in the amber inclusion. To investigate this possibility, we created surrogate datasets by artificially swapping sex identities and compared them with the original datasets. Because the female is always the leader in *C. formosanus*, the male is always the leader in the surrogate datasets. If the fossilized tandem was similar to the original datasets, then the female would be estimated as the tandem leader in *E. affinis*; if it were similar to the surrogate datasets, the male would be the leader.

We performed a principal component analysis on the 15 distances measured for all datasets (original datasets + surrogate datasets + fossil data) using the R prcomp() function. The four first principal components explained 97% of the observed variance. We used linear discriminant analysis for these four components to infer the leader-follower role of the amber inclusion of *E. affinis*. All data analyses were performed using R v. 4.0.1 (R Core Team, 2020).

## Acknowledgments

We thank Anna Prokhorova for providing ecological reconstruction art work; Jonas Damzen (www.amberinclusions.eu) for sharing the collection locality of the fossil; Kensei Kikuchi for his help with termite collection; and OIST imaging section for providing access to the microtomographic instrumentations. This study is supported by a JSPS Research Fellowships for Young Scientists CPD (20J00660) and a Grant-in-Aid for Early-Career Scientists (21K15168) to NM, and OIST core funding.

## Data accessibility

All data supporting this study’s findings, including µCT data in DICOM format and movement trajectories, and source codes for analyzing them are available at Github: github.com/nobuaki-mzmt/tandem-fossil. Data associated with the accepted version will be made available at Zenodo. The amber fossil was deposited at the National Museum, Prague (Czechia), as specimen NMP T3532.

## Author contributions

N.M.: conceptualization, data curation, formal analysis, funding acquisition, investigation, methodology, project administration, resources, supervision, validation, visualization, writing-original draft, writing-review & editing

S.H.: investigation, writing-review & editing

M.S.E.: writing-review & editing

T.B.: funding acquisition, resources, writing-review & editing

A.B.: conceptualization, data curation, formal analysis, investigation, methodology, project administration, resources, supervision, validation, visualization, writing-original draft, writing-review & editing

## Notes

https://github.com/nobuaki-mzmt/tandem-fossil

### Competing Interest Statement

The authors have declared no competing interest.

