## Supplementary material for "Extinct and extant termites reveal the fidelity of behavior fossilization in amber": Fig. S1-7; Table S1-2

This file includes
Supporting text S1
Figure S1 to S7
Table S1 to S2
Legend for Video S1

### Supporting Information Text

#### Text S1. Posture analysis of regular tandem runs.

Because the original videos for the datasets used as regular tandem runs (Mizumoto and Bourguignon, 2022) are not in high enough resolution to extract the coordinates of each body part, we used different videos for the posture analysis of regular tandem runs. Alates of *C. formosanus* were collected using light trapping in Okinawa, Japan, in May 2022. After collection, we brought them back to the lab and used individuals that shed their wings for behavioral observations. Individuals were separated by sex and kept on a moistened filter paper until the experiments. All observations were made within 12 hours after collection. Each pair was introduced in a Petri dish (ø = 90 mm) with moistened plaster and was recorded with a video camera (HC-X1500-K, Panasonic) for 15 minutes. All individuals were marked with one colored dot of paint (PX-20; Mitsubishi) on the abdomen to distinguish individual identity. In total, we obtained videos for 10 different pairs.

Instead of DeepLabCut (version 2.2.1.1), we used SLEAP (version 1.2.9) for body part tracking of regular tandem runs (Pereira et al., 2022). This is because our preliminary analysis found that DeepLabCut (or maDeepLabCut) did not successfully track regular tandem runs, while the SLEAP accurately tracked body parts of individuals during tandem runs. Seventeen body parts were labeled to form a skeleton for pose estimation: head, pronotum front, pronotum end, abdomen front, abdomen end, tips of antennae (left and right), middle points of antennae (left and right), bases of antennae (left and right), and distal ends of legs (six in total). For further analysis, we used data of three different points (head, pronotum, and abdomen tip). Note that we labeled the front tip of the head, instead of the middle of the head in SLEAP. We calculated the coordinate of the middle of the head from the coordinates of the front tip of the head and the front tip of the pronotum. We labeled 131 frames in total. To infer termite tracks across frames, top-down and centroid networks were trained within unet (interpolate -end was used as an anchor). Augmentation was performed only by rotating frames (from -180 to 180 degrees). Tracking was performed using the “simple” option with Intersection over Union (IOU) similarity method and Hungarian matching method. Misdetection of body parts was covered by interpolating the outputs with linear methods.

After obtaining three body part coordinates for each sex, we calculated the 15 pairwise distances and performed PCA with swapped datasets and all other datasets as described in the Methods section. The results show that the postures of regular tandems fell well within the region (i) and (v) in Fig. 3B (Fig. S5).

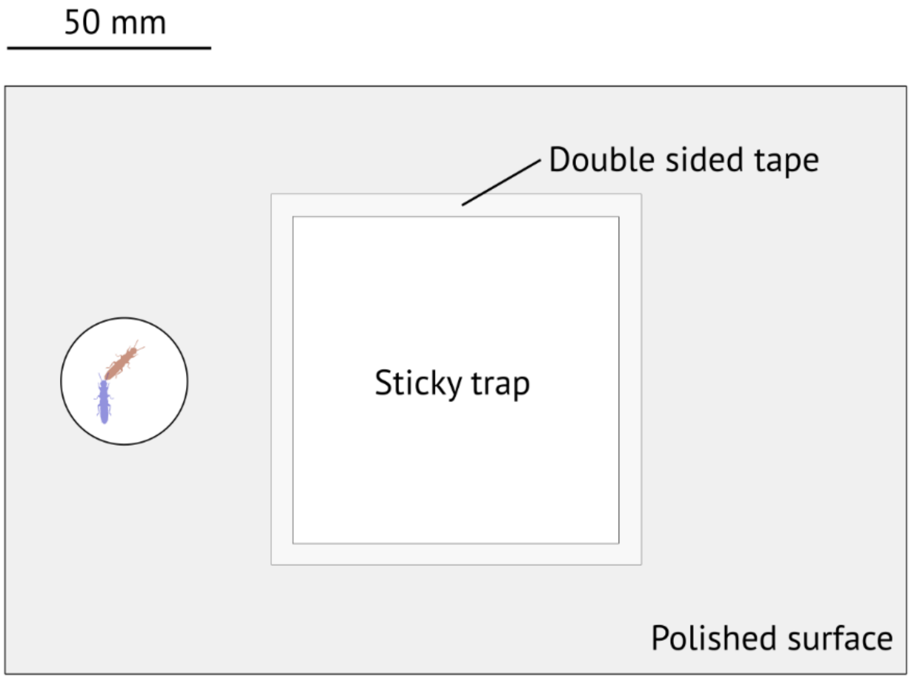


**Figure S1.** Experimental setup for observing tandem pair behavior on sticky surface simulating resin. A pair was isolated below a Petri dish (Φ=40mm) until a tandem run was initiated.

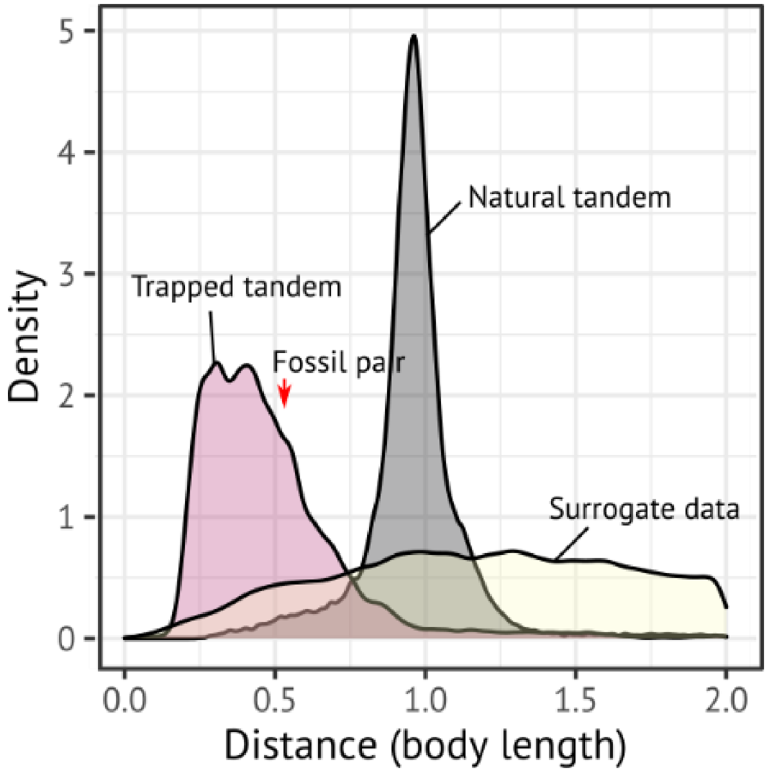


**Figure S2.** Distribution of distances between centroid of females and males across different contexts of tandem runs. The red arrow indicates the female-male distance in the amber inclusion.

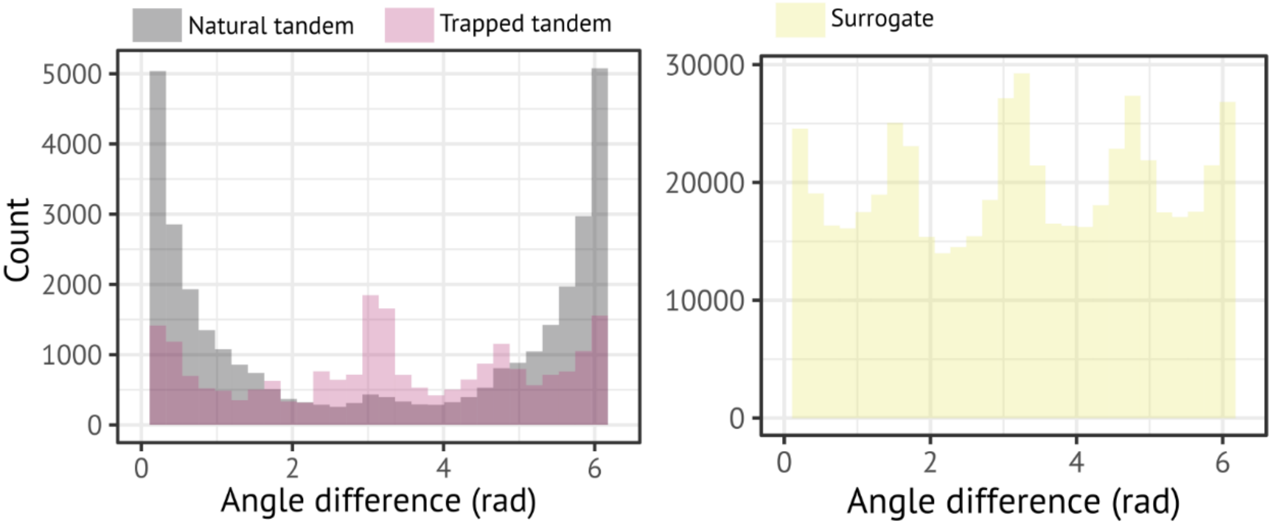


**Figure S3.** Distribution of the distance in heading direction between females and males across different context of tandem runs.

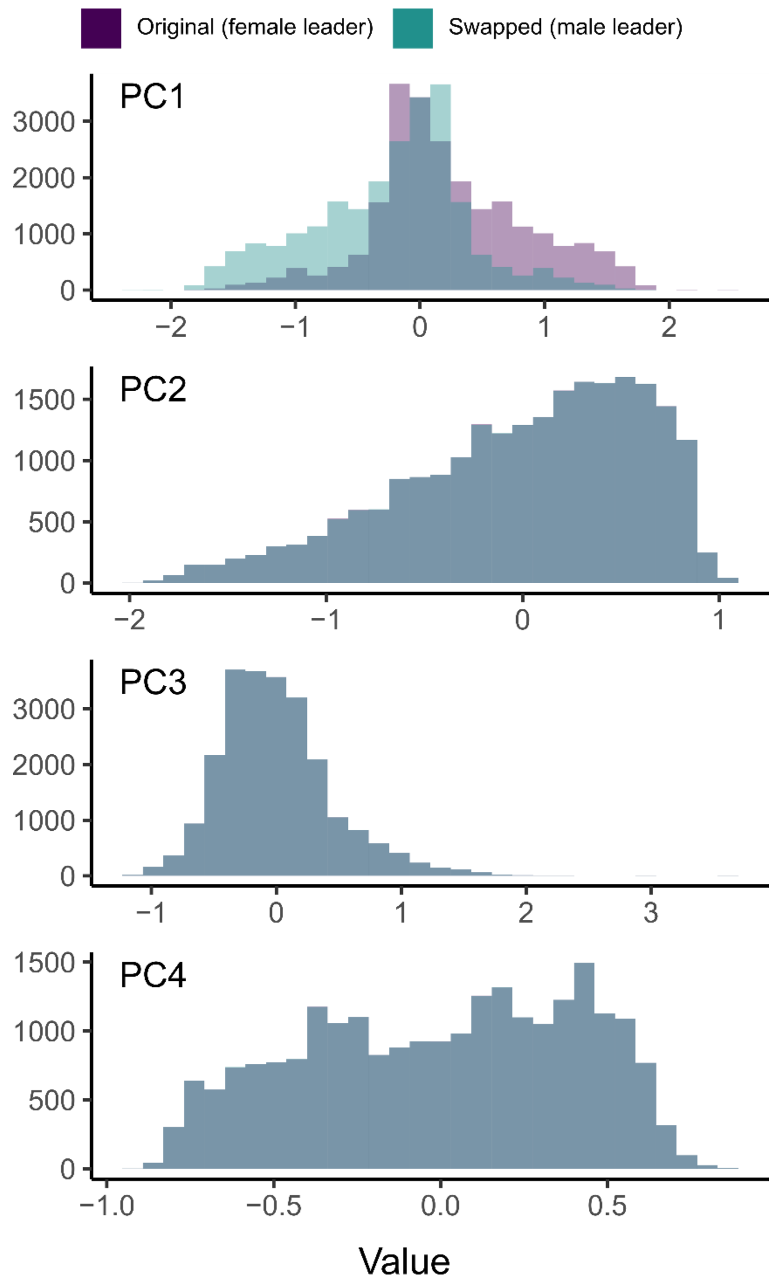


**Figure S4.** Distributions of values for PC1-PC4 in relation to the original or surrogate datasets. The four first principal components explained 97% of the observed variance. Among them, only the PC1 was asymmetric between the original and surrogate datasets, suggesting that the PC1 reflects the properties of postures related to the leader-follower roles.

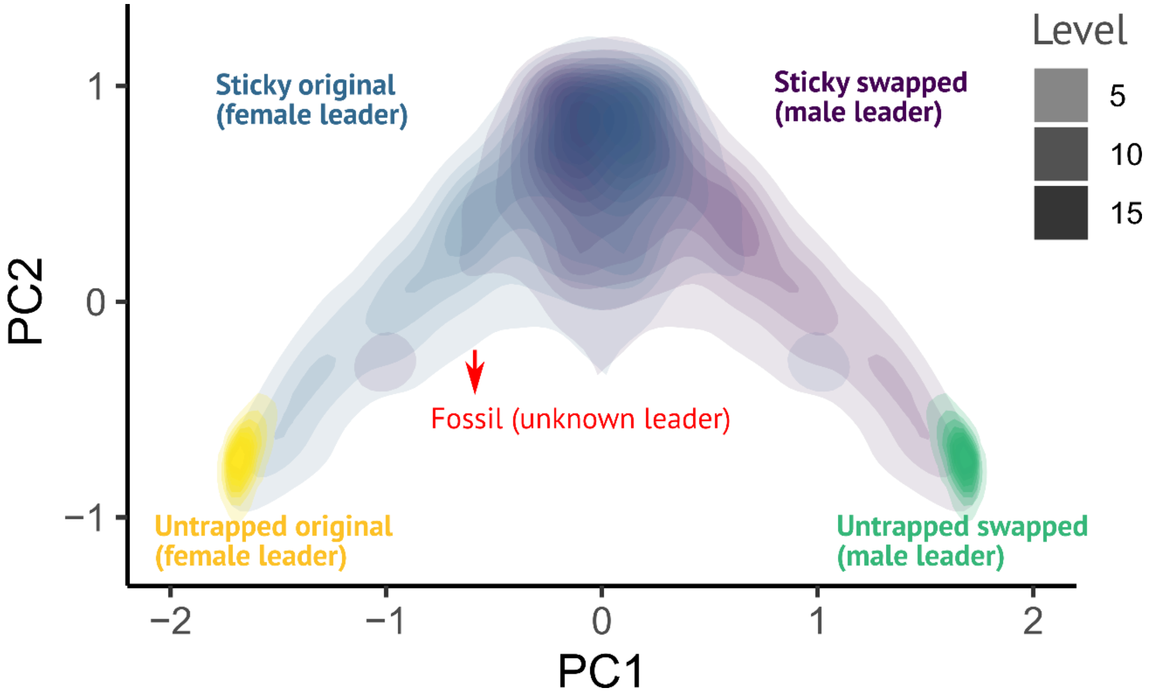


**Figure S5.** PCA results including posture datasets of untrapped tandem runs. See Text S1 for the methods.

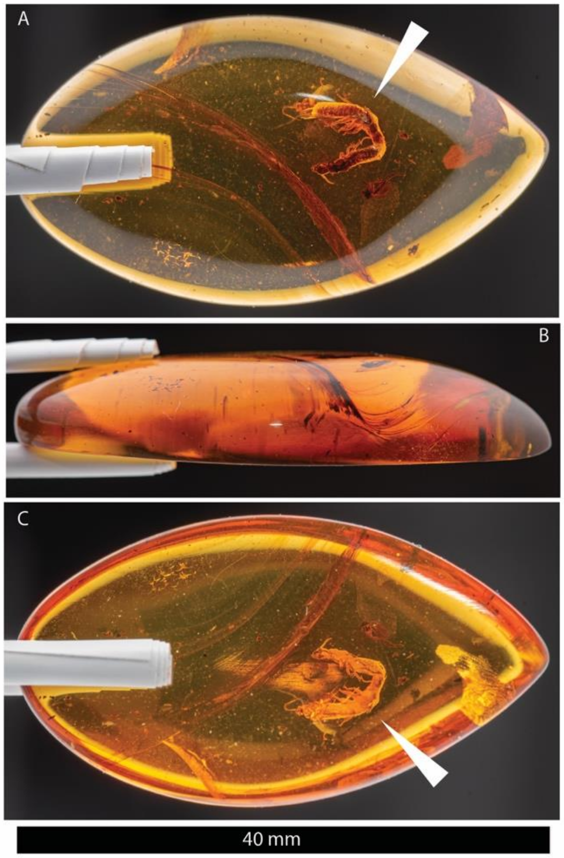


**Figure S6.** Overall habitus of the fossil tandem NMP T3532. (A) Dorsal side of the termite tandem; (B) sagittal view; and (C) ventral side of the termite tandem. White arrowheads indicate the position of the termite tandem.

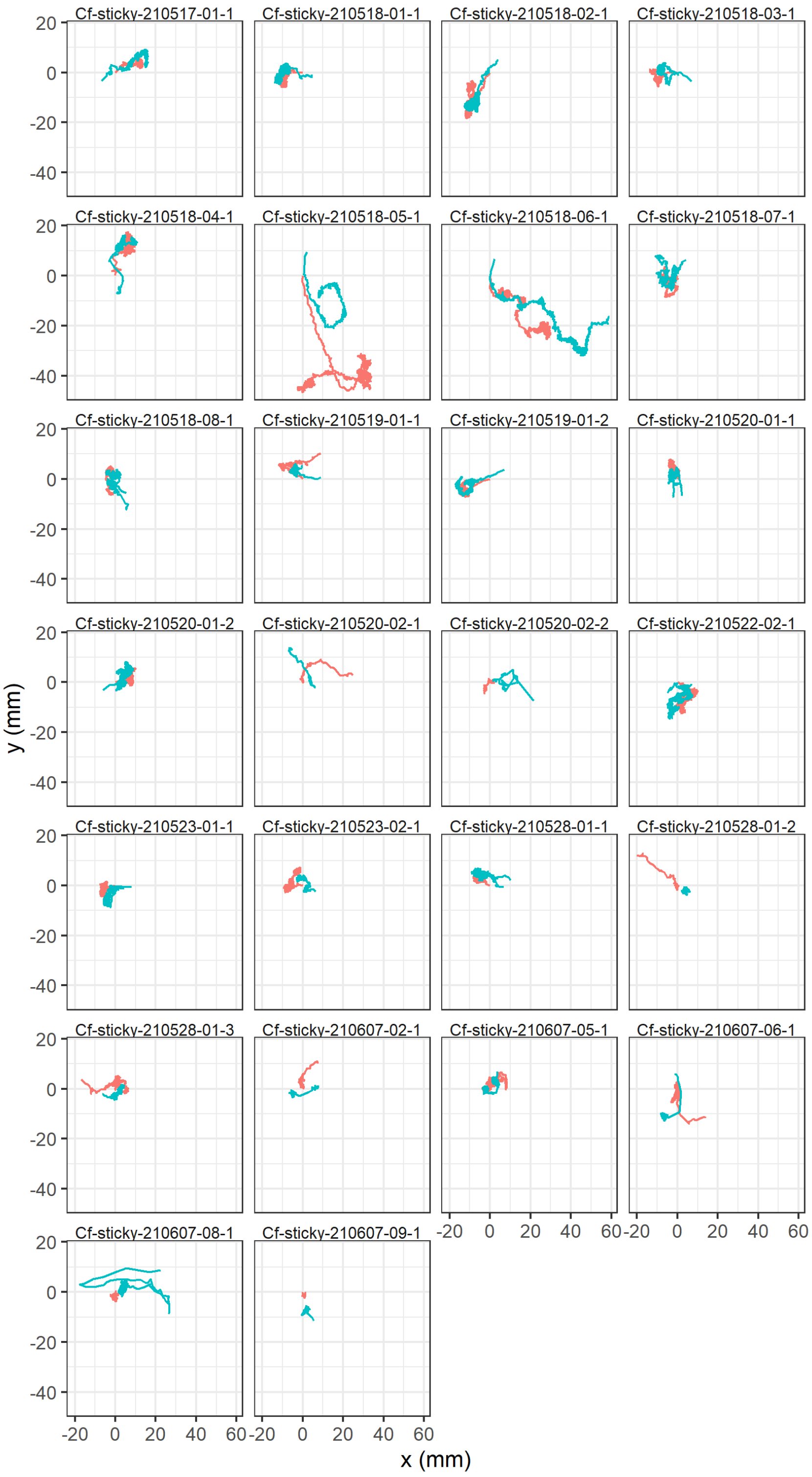


**Figure S7.** Trajectories of all pairs that entered the sticky trap area.

**Table S1.** Principle component loadings of all 15 pairwise distances used for posture PCA.

| Variables (distance between) | PC1 | PC2 |
|---|---|---|
| fhead_fpron | -0.0011 | -0.0053 |
| fhead_ftip | 0.0084 | -0.0149 |
| fhead_mhead | 0.0000 | -0.5303 |
| fhead_mpron | 0.0649 | -0.4983 |
| fhead_mtip | 0.5202 | -0.0172 |
| fpron_ftip | 0.0095 | -0.0080 |
| fpron_mhead | -0.0650 | -0.4983 |
| fpron_mpron | 0.0000 | -0.4684 |
| fpron_mtip | 0.4744 | -0.0107 |
| ftip_mhead | -0.5202 | -0.0171 |
| ftip_mpron | -0.4744 | -0.0106 |
| ftip_mtip | 0.0000 | 0.0370 |
| mhead_mpron | 0.0011 | -0.0053 |
| mhead_mtip | -0.0084 | -0.0149 |
| mpron_mtip | -0.0095 | -0.0080 |

fhead: female head, fpron: female pronotum, ftip: female abdomen tip, mfead: male head, mpron: male pronotum, mtip: male abdomen tip.

**Table S2.** Overview of scanning parameters.

| ScanID | Description | Voxel size ($\mu$m) | Source setting | Optical magnification | Source distance; detector distance (mm) | Exposure time (s) |
|---|---|---|---|---|---|---|
| fossil2b | abdominal tip of male and head of female | 3.45 | 40kV, 75$\mu$A | 4x | 27; 26 | 15 |
| fossil2c | abdominal tip of female | 2.17 | 40kV, 75$\mu$A | 4x | 26;55 | 17 |
| fossil2e | both termite individuals | 9.46 | 40kV, 75$\mu$A | 0.4x | 22;137 | 18 |

**Video S1.** Example of a termite mating pair caught by a sticky trap.
